# Optimized qRT-PCR approach for the detection of intra- and extra-cellular SARS-CoV-2 RNAs

**DOI:** 10.1101/2020.04.20.052258

**Authors:** Tuna Toptan, Sebastian Hoehl, Sandra Westhaus, Denisa Bojkova, Annemarie Berger, Björn Rotter, Klaus Hoffmeier, Sandra Ciesek, Marek Widera

## Abstract

The novel coronavirus SARS-CoV-2 is the causative agent of the acute respiratory disease COVID-19 which has become a global concern due to its rapid spread. Meanwhile, increased demand in testing has led to shortage of reagents, supplies, and compromised the performance of diagnostic laboratories in many countries. Both the world health organization (WHO) and the Center for Disease Control and Prevention (CDC) recommend multi-step RT-PCR assays using multiple primer and probe pairs, which might complicate interpretation of the test results especially for borderline cases. In this study, we describe an alternative RT-PCR approach for the detection of SARS-CoV-2 RNA that can be used for the probe-based detection of clinical isolates in the diagnostics as well as in research labs using a low cost SYBR green method. For the evaluation, we used samples from patients with confirmed SARS-CoV-2 infection and performed RT-PCR assays along with successive dilutions of RNA standards to determine the limit of detection. We identified an M-gene binding primer and probe pair highly suitable for quantitative detection of SARS-CoV-2 RNA for diagnostic and research purposes.

## 1. Introduction

In early January 2020, the novel severe respiratory coronavirus 2 (SARS-CoV-2) was discovered to be the causative agent of the coronavirus disease 2019 (COVID-19). SARS-CoV-2 outbreak was first detected in late December in Wuhan, China (Zhu et al., 2020) and was recently declared a pandemic by WHO (WHO, COVID-19 Situation Report – 51, published March 11^th^, (2020). We also observed a sharp increase of cases in Germany and containment measures were imposed to slow the progression. Detecting cases is difficult, as the disease is both very contagious, and can be clinically unremarkable in individuals shedding the virus (Hoehl et al., 2020; Rothe et al., 2020). Therefore, testing of suspected cases is of central importance but it is also required to guide patient care, and to apply appropriate hygiene measures in the hospitals to restrict nosocomial spread among patients and medical personnel (Klompas, 2020). The pandemic however, poses unprecedented challenges for institutions conducting the virologic testing, as measures of public health as well as patient care depend on timely, reliable results.

Quantitative reverse transcriptase polymerase chain reaction (qRT-PCR) is the gold standard in the detection of SARS-CoV-2. Distinct qRT-PCR testing protocols were swiftly established and made publicly available by the WHO (Corman et al., 2020), and by the Center for Disease Control (CDC) (CDC, 2020).

The protocol published by Corman et al. was designed before virus isolates were available and recommends a two-step process (Corman et al., 2020). In this workflow, the initial screening test is conducted with non-SARS-CoV-2-specific primers binding in the E-gene region coding for the Envelope small membrane protein. The confirmation of the E-gene positive samples should then be re-tested using primer pairs binding to the corona virus RNA dependent RNA polymerase (RdRP), which is specific in combination with a probe (P2) but less sensitive (Konrad et al., 2020). Here, the specificity of the confirmatory test relies on the probe target sequence, which has two mismatches compared to SARS-CoV and contains two wobble positions most likely hampering the sensitivity (**Table 2**). The CDC operates with a distinct protocol targeting three different regions within the N-gene encoding for the viral Nucleoprotein (CDC, 2020). In both protocols, multiple probes and primers are used in a multi-step PCR workflow, which is laborious and might also complicate interpretation of the results. In light of the current shortage of reagents, personnel, and equipment, a one-step PCR protocol achieving both high sensitivity and specificity would be beneficial for facing the SARS-CoV-2 pandemic. Optimized methods for both diagnostic and preemptive testing are essential to prevent the worst-case situation.

In this study, we designed an alternative PCR approach specific for the detection of SARS-CoV-2 RNA with primers binding in the M-gene encoding for the viral Membrane protein. This novel primer pair exerts a higher specificity than the E-gene protocol, but has significantly better sensitivity when compared with the RdRP-based PCR. As our approach requires fewer reagents and less-hands on time, it can be advantageous for early virus detection.

## 2. Results

### 2.1. Evaluation of specific PCR approaches for the detection of SARS-CoV-2 RNA

In several laboratories, non-specific PCR products in both SARS-CoV-2 negative patient samples and in the non-template control (NTC) using the WHO recommended SARS-CoV E-gene specific PCR (Corman et al., 2020) have been reported (Konrad et al., 2020) (personal communication). In order to confirm and further characterize these non-specific products in negative samples, we carried out additional PCR-based analyses. To this end, RNA from pharyngeal swab samples (**Table 1**) was isolated and subjected to WHO recommended qRT-PCR analysis using E (**Figure 1a**) and RdRP (**Figure 1b**) gene specific primers (**Table 2**). As positive controls we used RNA extracted from pharyngeal swabs of one asymptomatic, one mildly symptomatic passenger returning from Wuhan after initial quarantine (**Figure 1, Table 1**), (Hoehl et al., 2020). Both viral samples were passaged on Caco2 cells as described previously (Bojkova et al., 2020) and the resulting virus containing supernatants (FFM1 and 2) were used for subsequent analysis.

**Figure 1.**
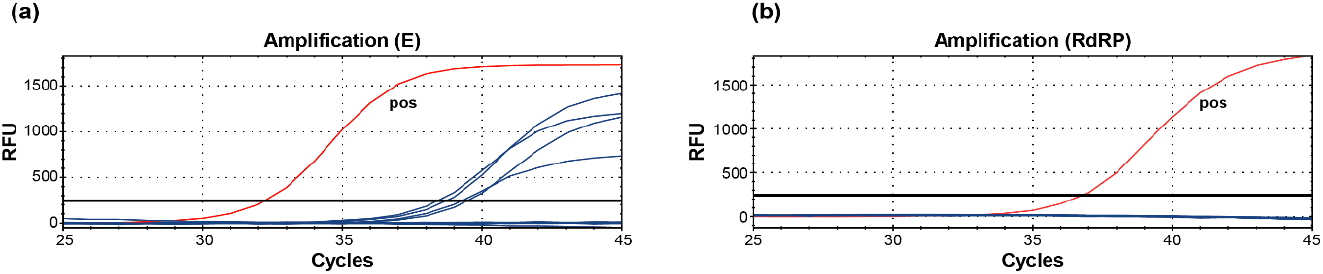
The WHO E-gene PCR for the pre-screening of SARS-CoV-2 using IVD-certified test kits (Roche) produces unspecific template-independent side-products. (**a**) Detection of SARS-CoV-2 false-positive samples tested with primers targeting for E-gene. (**b**) Confirmatory PCR using SARS-CoV-2 RdRP-specific primer and probe. RFU, relative fluorescence units; *pos,* positive control SARS-CoV-2 isolate FFM1.

**Table 1.**
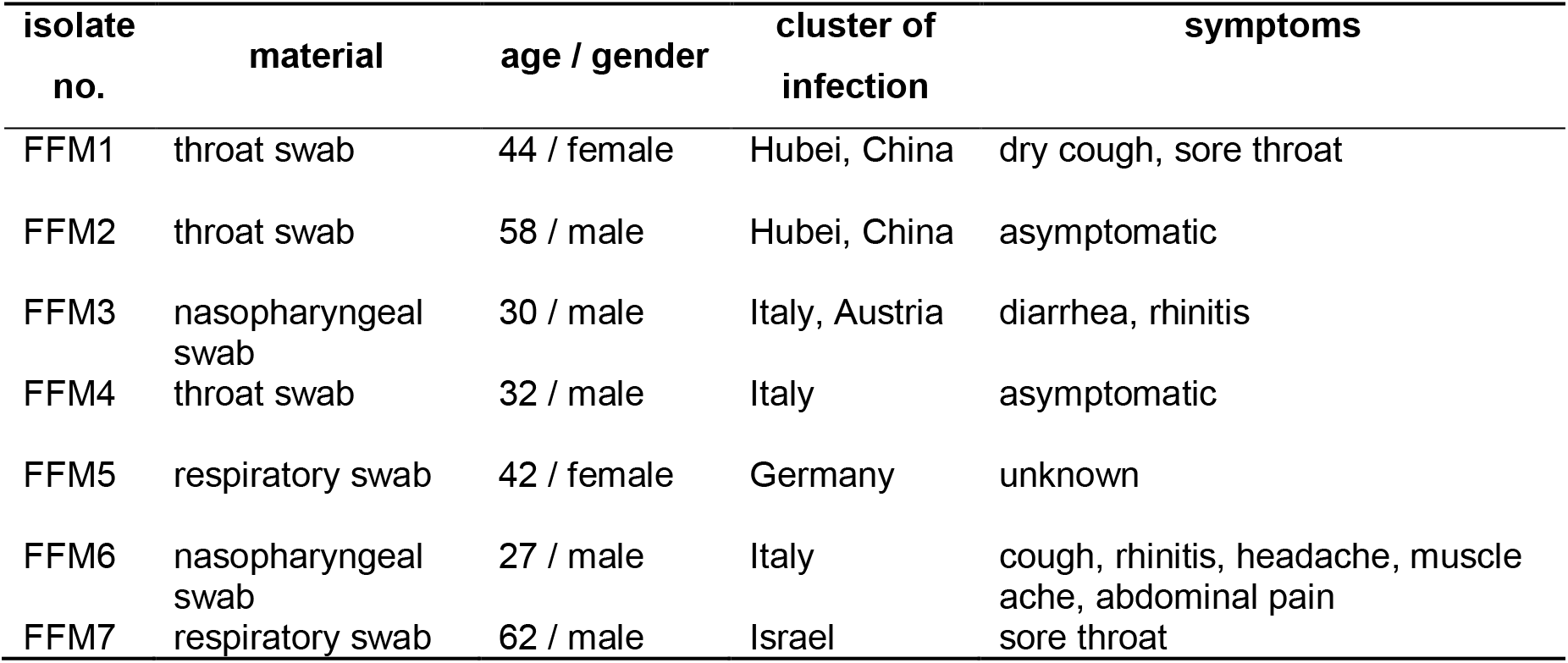
Compilation of patient samples and isolated viral strains in this study.

| isolate no. | material | age / gender | cluster of infection | symptoms |
| --- | --- | --- | --- | --- |
| FFM1 | throat swab | 44 / female | Hubei, China | dry cough, sore throat |
| FFM2 | throat swab | 58 / male | Hubei, China | asymptomatic |
| FFM3 | nasopharyngeal swab | 30 / male | Italy, Austria | diarrhea, rhinitis |
| FFM4 | throat swab | 32 / male | Italy | asymptomatic |
| FFM5 | respiratory swab | 42 / female | Germany | unknown |
| FFM6 | nasopharyngeal swab | 27 / male | Italy | cough, rhinitis, headache, muscle ache, abdominal pain |
| FFM7 | respiratory swab | 62 / male | Israel | sore throat |

**Table 2.**
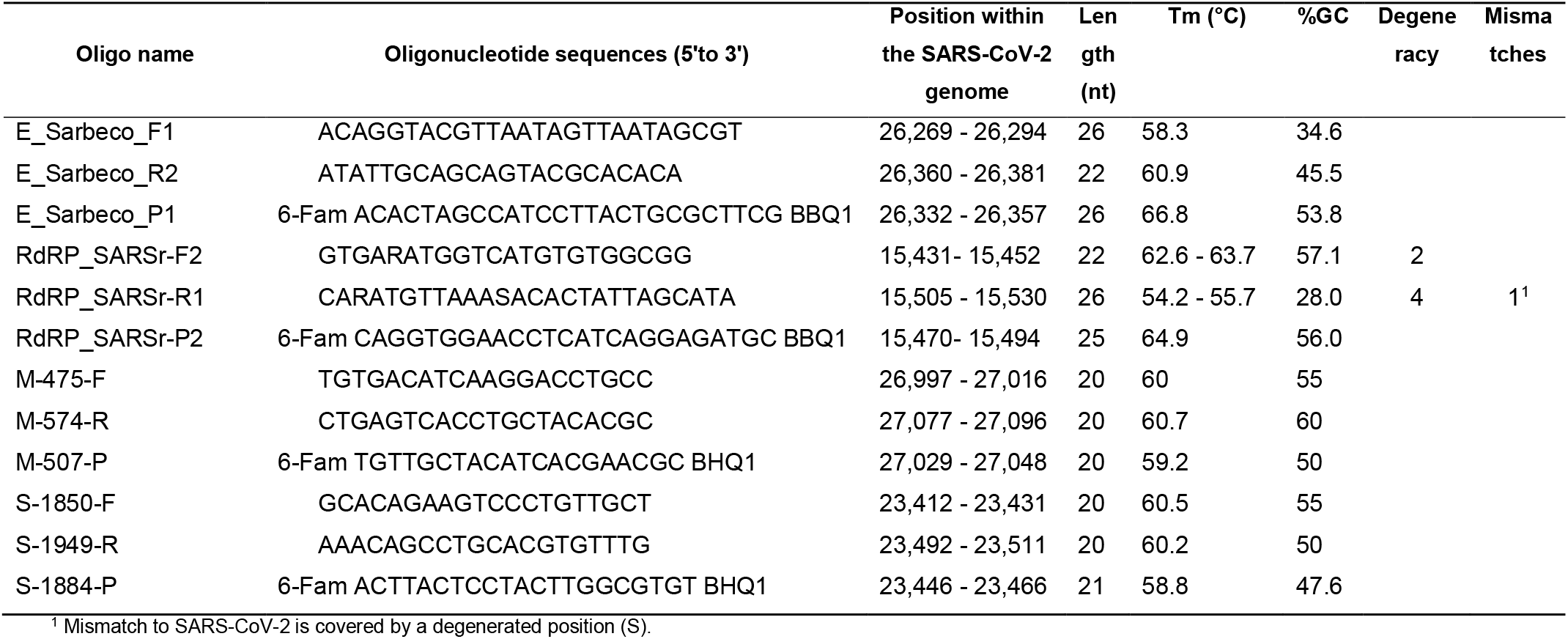
Oligonucleotides used for the detection of SARS-CoV-2 RNA. IUPAC codes: S = G or C; R = A or G.:

Four out of 48 (8.33%) tested swab samples showed a positive signal in the E-gene PCR (**Figure 1a**), which were of the same size as the specific amplicons (data not shown). However, the confirmatory RdRP-gene PCR was negative for all patient samples except the positive control (**Figure 1b**). This data indicates that using the two-step WHO PCR protocol might complicate interpretation of the test results especially for cases with higher Cq-values.

Since the RdRP-gene PCR is more specific than the E-gene PCR but is also described as less sensitive (Konrad et al., 2020), we aimed to develop a novel approach using specifically designed primer and probe pairs. For optimal primer design consensus sequences were aligned to the reference SARS-CoV-2 genome sequence and analyzed for primers binding in E, N, Orf1, M, and S regions (**Supplementary Figure 1**) that would allow SARS-CoV-2 but not SARS-CoV amplification. For experimental confirmation, we used human SARS-CoV strain Frankfurt 1 (NC_004718) and SARS2-CoV-2 RNA samples (FFM1 and FFM2) and compared the PCR performances using serial dilutions (**Supplementary Figure 1a**). Due to limited linearity, E and N gene specific primers were excluded from further analysis while S and M gene based PCRs (**Table 1**), which were more sensitive than Orf-1 (**Supplementary Figure 1b**), proved to be suitable for linear SARS-CoV-2 RNA detection including samples with low viral loads (~Cq 40) (**Supplementary Figure 1c**). Since M gene PCR was superior by approximately Cq 1.73 +/− 0.26 earlier detection (Cq 1.48 +/− 0.15 for SARS-CoV, **Supplementary Figure 1d**), we continued with M gene PCR and further characterized the limit of detection.

Using plasmid DNA constructs that harbor the conserved SARS-CoV-2 amplicon sequences, we generated standard curves and compared the M-gene PCR (**Figure 2b**) with the established WHO RdRP-gene PCR approach (**Figure 2a**). Using the same dilution series of plasmid samples, the M gene based method allowed earlier detection (**Figure 2c**). To determine the analytical sensitivity of the M-gene based approach, we additionally used *in vitro*-transcribed RNA standards and tested four replicates to determine the limit of detection (**Figure 2d-f**). We were able to detect a single RNA copy per reaction by M-Gene qRT-PCR. SYBR-green based melting curve analysis revealed a melting point at 80°C for all samples (**Supplementary Figure 2**). To further characterize the capacity of M-gene PCR to detect high viral loads, we performed SYBR-green based PCR and quantified high loads of virus cell culture supernatants (**Supplementary Figure 2d)**.

**Figure 2:**
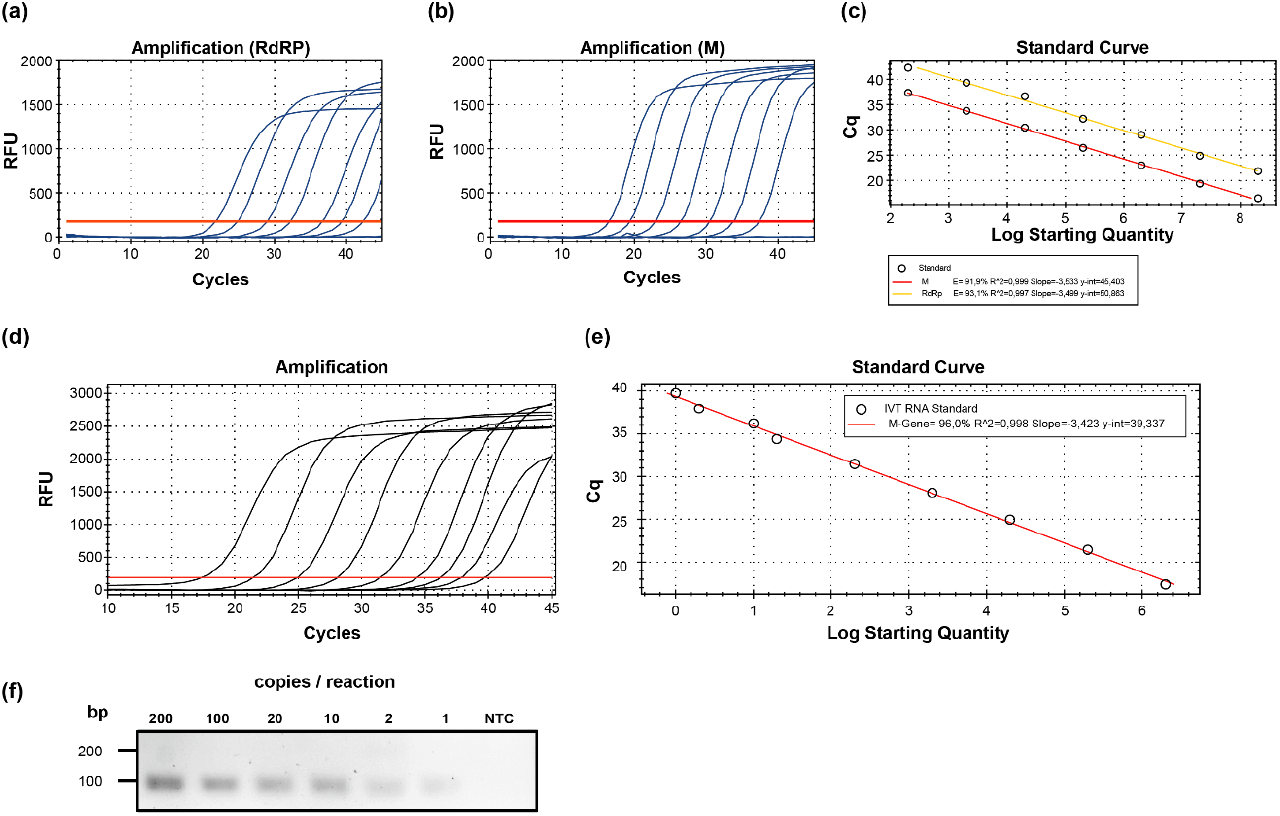
Characteristics of M-gene based qRT-PCR. (**a**) Representative amplification curves of WHO RdRP- and (**b**) SARS-CoV-2 M gene specific qRT-PCR using plasmid standards copy numbers. (**c**) Standard curves of both genes determined with plasmid templates. The Cq was plotted against the log starting quantity of the indicated plasmid DNA template. (**d**) Representative amplification curves and (**e**) standard curves of SARS-CoV-2 M gene specific qRT-PCR using *in vitro* transcribed RNA templates. Log starting quantity (copies/reaction) is indicated and plotted against the Cq. (**f**) M-gene amplicons visualized in a 2% agarose gel. The RNA copy numbers are shown on the top. RFU, relative fluorescence units; bp, base pairs; Cq, quantification cycle, NTC, no template control (H2O); E, amplification efficiency.

In order to confirm that the M-gene specific PCR primers and probes cover all known isolates and thus enable detection, an alignment of published SARS-CoV-2 full-length isolates known at the time of submission (n = 165) and viral isolates sequenced during this study was carried out (**Figure 3, Supplementary Figure 3**). Both forward and reverse primers had a 100% identity and only one isolate of 165 (0.006%) had a mismatch to the consensus sequence C/T) at position 18 of the DNA probe (**Figure 3**). However, this mismatch would not affect the detection, but possibly reduce the sensitivity but not prevent binding of the probe.

**Figure 3:**
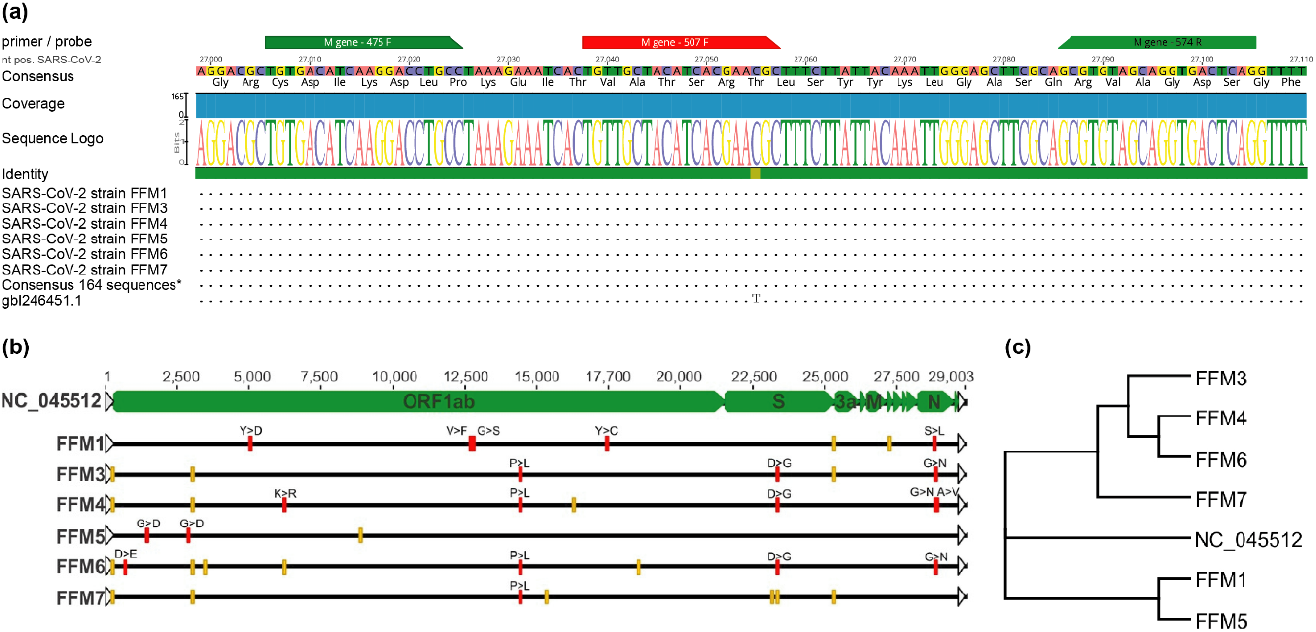
SARS-CoV-2 sequences are highly conserved in M-gene. (**a**) Primer binding sites of M-gene primers in a partial alignment of SARS-CoV-2 related sequences (n =165) showing oligonucleotide relevant binding regions. Graphical representation of the sequence conservation of nucleotides and amino acid sequences are depicted as sequence logo. The relative sizes of the letters illustrate their frequency in the compared sequences. The height of the letters is a measure of the information content of the position in bits. (**b**) Multiple sequence alignments of the SARS-CoV-2 strains described in this study as sequenced. Silent mutations are indicated in yellow boxes, amino acid substitutions are shown as red boxes using the single amino acid code. (**c**) Phylogenetic tree of SARS-CoV-2 isolates used in this study. SARS-CoV-2 reference genome NC_0445512. Phylogenetic tree analysis of SARS-CoV-2 isolates.

In conclusion, our M-gene based qRT-PCR detection of SARS-CoV-2 RNA was at least as specific as the RdRP PCR recommended for confirmation by the WHO, but showed a significantly higher sensitivity. Importantly, unspecific signals as observed in the E-gen PCR were not detected.

### 2.3. Detection of SARS-CoV-2 in clinical and research samples using E-, RdRP-, and M-gene specific protocols

In order to validate our method, we re-tested clinically relevant samples that have been qualitatively tested positive for SARS-CoV-2 RNA during routine diagnostics (**Supplementary Table 1**). WHO recommended RdRP primer pairs were used for confirmation. As negative controls we included 8 negative samples (**Figure 1, Supplementary Table 1)**. As described above, unspecific E-gene amplicons were detected in a test kit specific manner possibly due to the reagents used (Konrad et al., 2020). Therefore, we additionally compared the performances of two research kits (New England Biolabs) and one *in vitro* diagnostic (IVD) certified test kit (Roche Viral Multiplex RNA Kit) using M-gene and RdRP-gene primers. Overall, we observed lower Cq values with all three kits for M-gene when compared to RdRP-gene PCR (**Figure 4**). Using the Luna OneStep Probe kit (Luna Universal Probe One-Step RT-qPCR Kit, NEB) M-gene PCR was significantly more sensitive among all tested kits with a difference of approx. 10 Cq values Luna Universal Probe One-Step RT-qPCR Kit in comparison to the RdRP specific primer pairs. With the SYBR green based kit (Luna Universal One-Step RT-qPCR Kit) and Roche IVT kit (LightCycler^®^ Multiplex RNA Virus Master) we observed four and one Cq value differences respectively, verifying that detection of M gene is superior to RdRP-gene PCR. To further evaluate whether the newly developed method is also suitable for detecting intracellular virus RNA, we infected Vero and Caco2 cells with SARS-CoV-2 strain FFM1. Using M-gene PCR we were able to detect very low copy numbers of viral RNA in infected Vero cells, while RdRP-gene PCR was limited in sensitivity (**Figure 5a**). Furthermore, we infected Caco2 cells and performed an intracellular replication curve and monitored viral replication and genome copy numbers, respectively. Viral RNA was detectable at the time points 3, 6, 12, and 24 h post infection in a linear fashion (**Figure 5b-c**).

**Figure 4:**
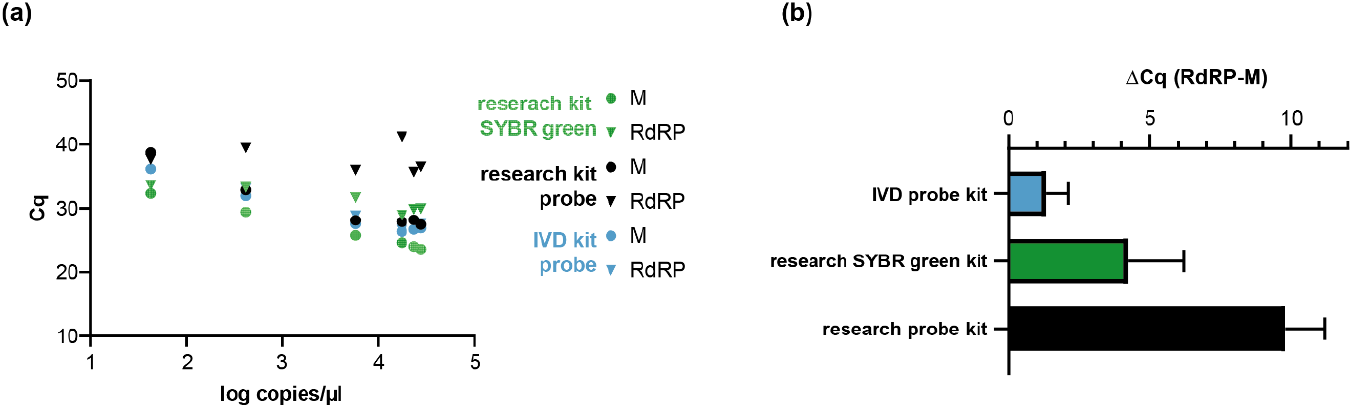
Patient sample detection using M versus RdRP specific primers. (**a**) Comparison of M and RdRP specific primer and probes measured with the indicated one step qRT-PCR kits. (**b**) Mean values of the Cq difference between RdRP and M gene PCR. Cq, quantification cycle.

**Figure 5:**
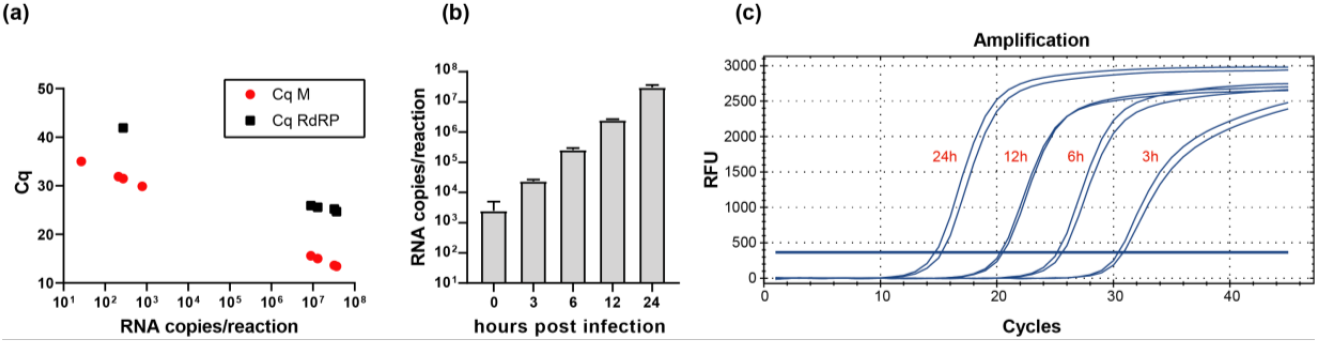
Detection of intracellular SARS-CoV-2 RNAs. (a) Vero cells were infected with SARS-CoV-2 strain FFM1 with high and low MOI and 24 h post infection the RNA was subjected to M and RdRP-Gene specific one step qRT-PCR-analysis. Intracellular copy numbers (**b**) and representative amplification curves (**c**) of SARS-CoV-2 strain FFM1 infected Caco2 cells. Total cellular RNA including viral RNA was harvested at the indicated time points and subjected to M-gene specific qRT-PCR analysis.

In conclusion, non-specific PCR products and limited sensitivity pose a major problem using the WHO protocol for SARS-CoV-2 detection and could lead to numerous unnecessary confirmation tests. Our newly developed PCR protocol is suitable to save time and resources since pre-screening is no longer necessary. Thus, this approach might be used as a cost-effective alternative to the E- and RdRP-based protocol.

## 3. Discussion

Countermeasures against COVID-19 depend on testing with the highest sensitivity and specificity possible. We and others (Konrad et al., 2020) (personal communication) determined certain drawbacks in the two-step PCR protocol recommended by the WHO as unspecific signals for E-gene PCR may arise due to combination of several factors including primer dimers, unspecific binding of the primers and probes, RT-PCR kit and thermocycler dependent differences. In commercially available kits, primers and probes, as well as the buffers and enzyme quantities and properties have been matched to one another to reduce unspecific amplification (Konrad et al., 2020). In times of increasing reagent shortages, however, a simple protocol should be available that can be quickly adapted with universal test kits (including research kits) and is easy to evaluate would be beneficial. In this study, we evaluated several primer and probe pairs designed *in silico* and identified M-gene specific primer and probe (**Table 2**) to be highly versatile for detection of SARS-CoV-2 RNA in a large selection of samples. We analyzed samples from patients presented with different symptoms, viral loads and degree of infection (**Table 1 and Supplementary Table 1**) and compared isolates with varying demographic characteristics originating from several infection clusters (**Table 1**). Interestingly, via sequence comparison based on the phylogenetic analysis we were able to show that isolate FFM5 with unknown origin shows a high degree of relationship to the FFM1 isolate from China indicating the same origin (**Figure 3**). Anyway, the SARS-CoV-2 M-gene PCR was able to detect SARS-CoV-2 from all clusters.

In a direct comparison of the RdRP PCR recommended by the WHO and the M-Gen PCR developed in this study, the M-Gen PCR was superior in terms of sensitivity using multiple test kits. Since we used primers that specifically bind SARS-CoV-2 RNA, this was to be expected. The reduced sensitivity of the WHO recommended RdRP-PCR may be explained by the presence of wobble pairs (**Table 2**) in the primer and probe sequences (Corman et al., 2020). Nevertheless, in a situation of crisis, where the availability of reagents, equipment as well as labor of qualified personnel are limited, a reliable and convenient one-step PCR protocol with optimum sensitivity would be beneficial.

Our comparison analysis for M- and RdRP-gene PCR were consistent and reproducible for detecting RNA from progeny virus particles and also intracellular viral mRNAs extracted from infected Caco2 and Vero cells even if some of the subgenomic viral mRNAs do not harbor the target sequence (Kim et al., 2020). We also have shown that the method is also suitable for inexpensive SYBR-green based PCR assays (**Supplementary figure 2d**). Thus, M gene PCR allows the quantification of very low and very high viral loads.

## 4. Materials and Methods

### Cell culture and virus preparation

Caco2 and Vero cells (African Green monkey kidney) were cultured in Minimum Essential Medium (MEM) supplemented with 10% fetal calf serum (FCS), 100 IU/ml of penicillin and 100 g/ml of streptomycin. SARS-CoV-2 isolate (Frankfurt 1 was obtained from throat swab of a patient diagnosed with SARS-CoV-2, hospitalized in the isolation unit of Frankfurt University Hospital (Germany) (Hoehl et al., 2020). SARS-CoV isolate FFM-1 was isolated from patient diagnosed with SARS-CoV infection in 2003 at Frankfurt University Hospital (Drosten et al., 2003). SARS-CoV-2 was propagated in Caco-2 cells using MEM, while SARS-CoV was grown in Vero cells both using with 1% FCS Virus stocks were stored at −80°C. Viral titers were determined by TCID_50_ or qRT-PCR. In accordance to the decision of the Committee on Biological Agents (ABAS) and Central Committee for Biological Safety (ZKBS) all work involving infectious SARS-CoV-2 and SARS-CoV was performed under biosafety level 3 (BSL-3) conditions in a BSL-3 facility. For intracellular RNA testing, 1x 10^5^ Vero cells were seeded per well in a 12-well plate. Cells were inoculated with SARS-CoV-2 or SARS-CoV suspension at an MOI of 0.1 for one hour at 37°C, 5% CO_2_. Subsequently cells were rinsed with PBS, replenished with fresh media and incubated for another 7 days before harvesting for RNA extraction.

For time point analysis, Caco2 cells were infected with SARS-CoV-2 (0.01 MOI) in MEM media supplemented with 1% FCS, at 37°C, 5% CO_2_. Cells were harvested for RNA extraction 0, 3, 6, 12 and 24 h post infection using TRI Reagent (Sigma). Following chloroform extraction and isopropanol precipitation, RNA was dissolved in nuclease free water and treated with DNase I (Qiagen) for 15 min at 37°C. DNase I treated RNA was subjected to RT-PCR analysis.

### Quantification of SARS-CoV-2 RNA

SARS-CoV-2 RNA from cell culture supernatant samples was isolated using AVL buffer and the QIAamp Viral RNA Kit (Qiagen) according to the manufacturer’s instructions. Intracellular RNAs were isolated using RNeasy Mini Kit (Qiagen) as described by the manufacturer. RNA was subjected to OneStep qRT-PCR analysis using Luna Universal One-Step RT-qPCR Kit (New England Biolabs) or Luna Universal Probe One-Step RT-qPCR Kit (New England Biolabs) or LightCycler^®^ Multiplex RNA Virus Master (Roche) using CFX96 Real-Time System, C1000 Touch Thermal Cycler. Primer pairs for E-, S- and M-gene specific PCRs were used in equimolar concentrations (0.4 μM each per reaction). For RdRP-primer pairs were used according to Corman et al. (Corman et al., 2020) with 0.6 μM and 0.8 μM for forward and reverse primers, respectively. The cycling conditions were used according to the manufacturer’s instructions. Briefly, for SYBR green and probe based Luna Universal One-Step RT-qPCR Kits, 2 μl of RNA were subjected to reverse transcription performed at 55°C for 10 minutes. Initial denaturation was 1 min at 95°C followed by 45 cycles of denaturation for 10 seconds, extension for 30 seconds at 60°C. Melt curve analysis (SYBR green) was performed from 65-95°C with an increment of 0.5°C each 5 seconds. For IVD approved LightCycler^®^ Multiplex RNA Virus Master (Roche) 5 μl of template RNA were used. Reverse transcription was performed at 55°C for 10 minutes. Initial denaturation was allowed for 30 seconds at 95°C followed by 45 cycles of denaturation for 5 seconds, extension for 30 seconds at 60°C and final cool-down to 40°C for 30 seconds. The PCR runs were analyzed with Bio-Rad CFX Manager software version 3.1 (Bio Rad Laboratories).

### Primer Design

For primers design and validation 165 available SARS-CoV-2 full-length sequences were aligned using NCBI Virus Variation Resource (Hatcher et al., 2017) and a consensus was made with 99% identity cutoff. Primer3 and Geneious Prime^®^ software version 2020.0.5 (Biomatters Ltd.) were used to design primer and probes matching the consensus sequence. Oligo characteristics were calculated using a modified version of Primer3 2.3.7. Positions within the SARS-CoV-2 genome according to accession number MN908947. Illustration were made with Geneious software.

### Generation of DNA and RNA standard curves

Standard curves were created using plasmid DNA harboring the corresponding amplicon target sequence according to GenBank Accession number NC_045512. For *in vitro* transcription, pCR2.1 based plasmid DNA (pCR2.1 SARS-CoV-2 M (475-574)) was linearized with BamHI. In vitro transcription was carried out using the HiScribeT7 High Yield kit (NEB) according to the manufacturer’s instructions. Absorbance-based measurement of the RNA yield was performed using the Genesys 10S UV-Vis Spectrophotometer (Thermo Scientific).

### Illumina NGS Sequencing of SARS-CoV-2 isolates

Caco2 cells were infected with different viral strains (FFM1-FFM7) at an MOI 0.01. Cell culture supernatant was harvested 48 h after infection, precleared at 2000xg for 10 min at room temperature. Virus particle containing supernatant (20 ml) was overlaid on 25% sucrose in PBS and centrifuged at 26,000 rpm for 3 h at 4C using SW28 rotor (Beckman Coulter). Pellets were dissolved in PBS overnight at 4C and 1/3 of used for RNA was extraction using Macherey Nagel Virus RNA isolation kit without carrier RNA according to the manufacturer’s recommendations. The libraries were prepared and sequenced by GenXPro GmbH, Frankfurt a. M., Germany. Briefly, quality was controlled on a LabChip GXII platform and RNA-seq libraries were prepared using the NEBNext Ultra II Directional RNA-Seq kit according to the manufacturers recommendations. Sequencing was performed on an Illumina NextSeq 500 platform using 75 cycles. Read mapping was performed using NC_045512 as a reference sequence. The viral nucleotide sequences are available at the following GenBank accession numbers: FFM1, MT358638; FFM3, MT358639; FFM4, MT358640; FFM5, MT358641; FFM6, MT358642; FFM7, MT358643.

## Acknowledgments

The authors thank Marhild Kortenbusch, Christiane Pallas and Lena Stegmann for excellent technical assistance. We thank the numerous donations and the support of SARS-CoV-2 research.

---

**Supplementary Figure 1.**
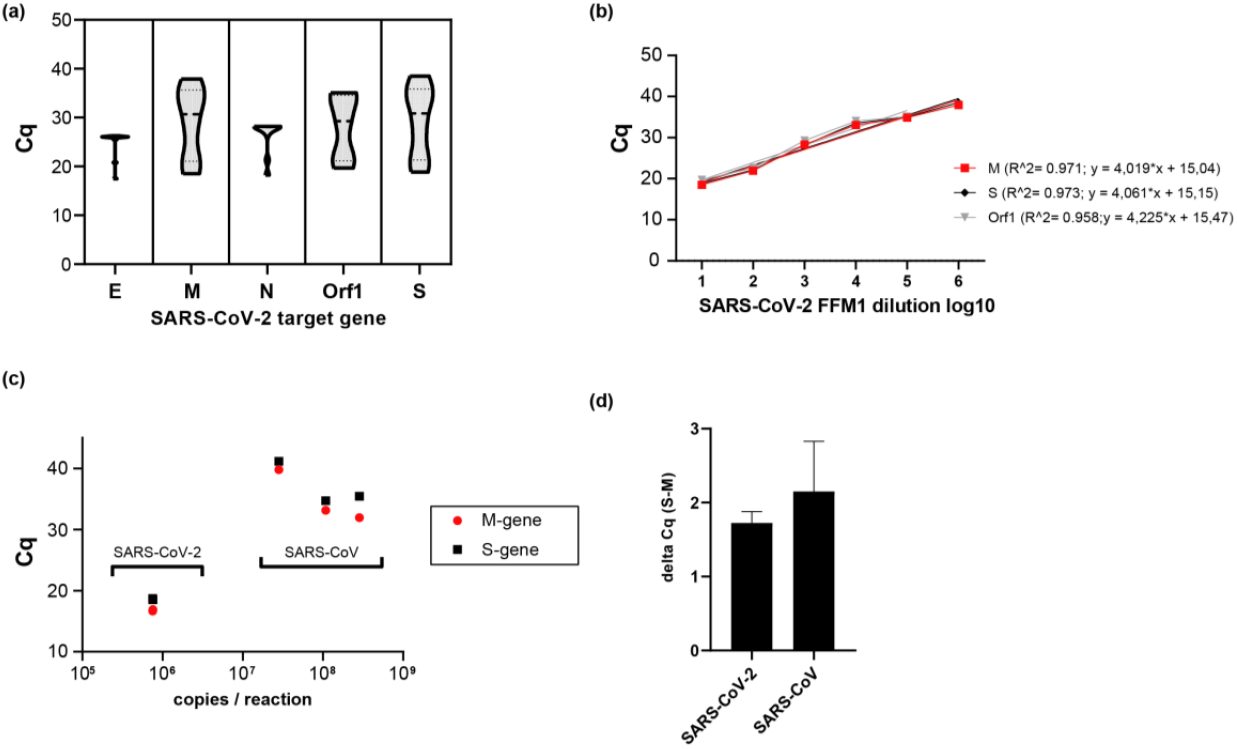
Evaluation of SARS-CoV-- gene specific qRT-PCR. (**a**) Violin plots showing Cq distributions of successive RNA dilutions of SARS-CoV-2 isolate FFM1. (**b**) Amplification curves of the dilution series were used for valuation of S-and M-gene specific qRT-PCR. (**a**) Detection of SARS-CoV-1 and SARS-COV-2 RNA samples (n=3) using M- and S-gene specific primer and probe pairs. (**b**) Linearity of M- and S-gene qRT-PCRs. (**c**) Detection of and SARS-CoV strain Frankfurt 1 (NC_004718) and SARS-CoV-2 strain FFM1 RNA using M and S-gene specific primer and probes. (**d**) Comparison of Cq values obtained wit S or M specific primes. RFU, relative fluorescence units; bp, base pairs; Cq, quantification cycle.

**Suppl. Figure 2:**
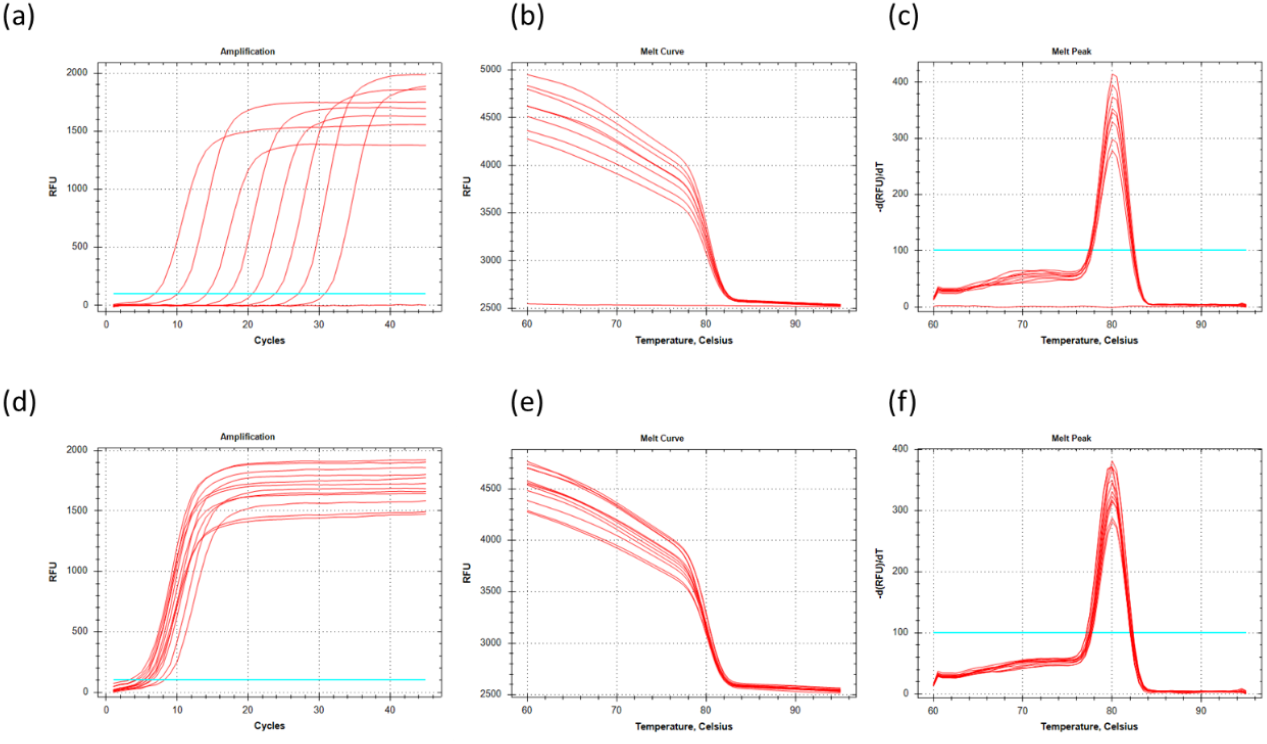
Detection of SARS-CoV-2 RNA with M-gene specific PCR primer and probes using SYBR green. (**a**) Amplification curve of *in vitro* transcribed RNA-standards and concentrated virus supernatants (**d**). (**b and e**) melt curve analysis and (**c and f**) melt peaks of M-gene amplicons. RFU, relative fluorescence units; bp, base pairs; Cq, quantification cycle.

**Suppl. Figure 3:**
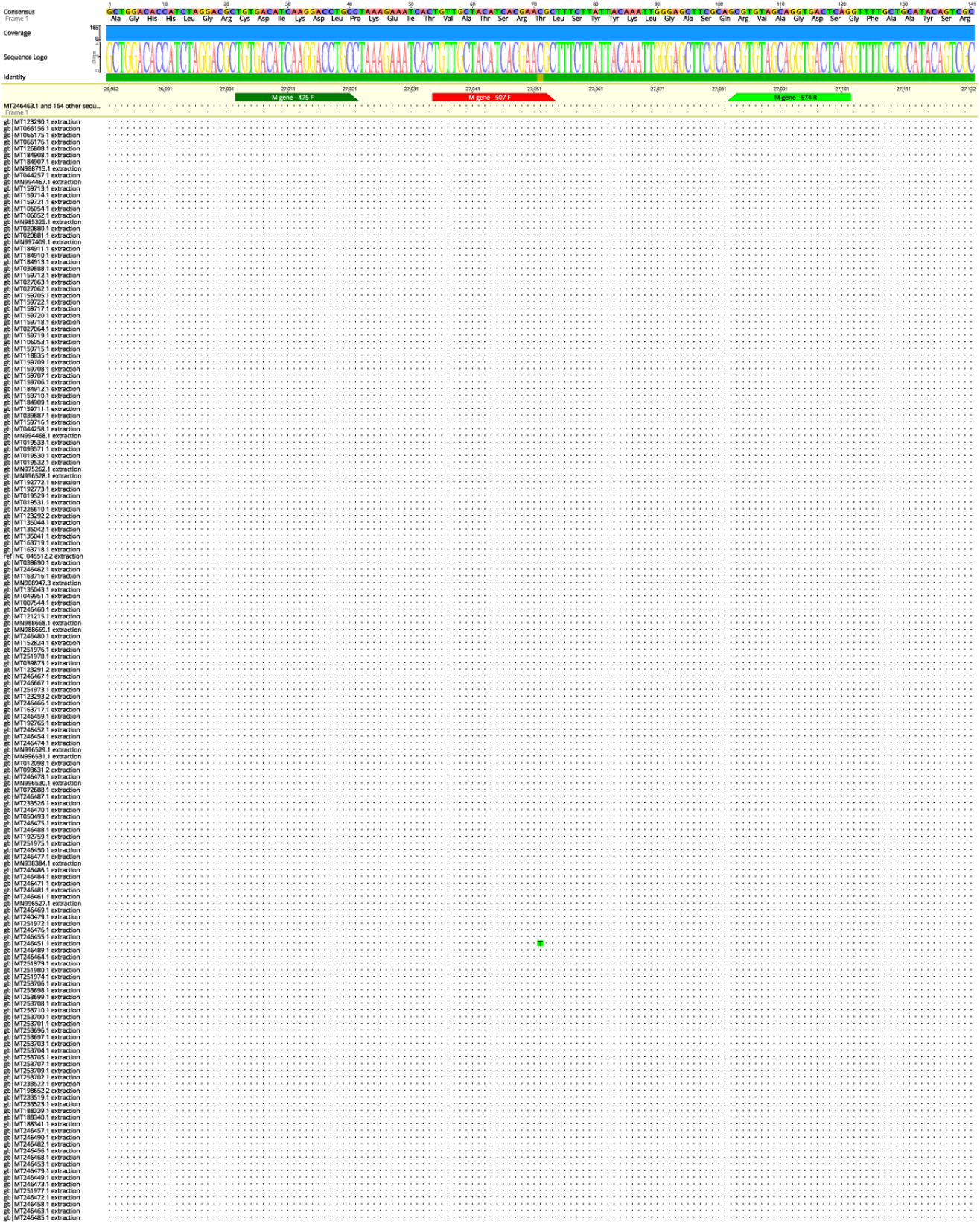
Sequence Alignment of 165 CoV-samples (extension to figure 3). Primer binding sites of M-gene primers are indicated. Graphical representation of the sequence conservation of nucleotides and amino acid sequences are depicted as sequence logo. The relative sizes of the letters illustrate their frequency in the compared sequences. The height of the letters is a measure of the information content of the position in bits.

**Supplementary Table 1.**
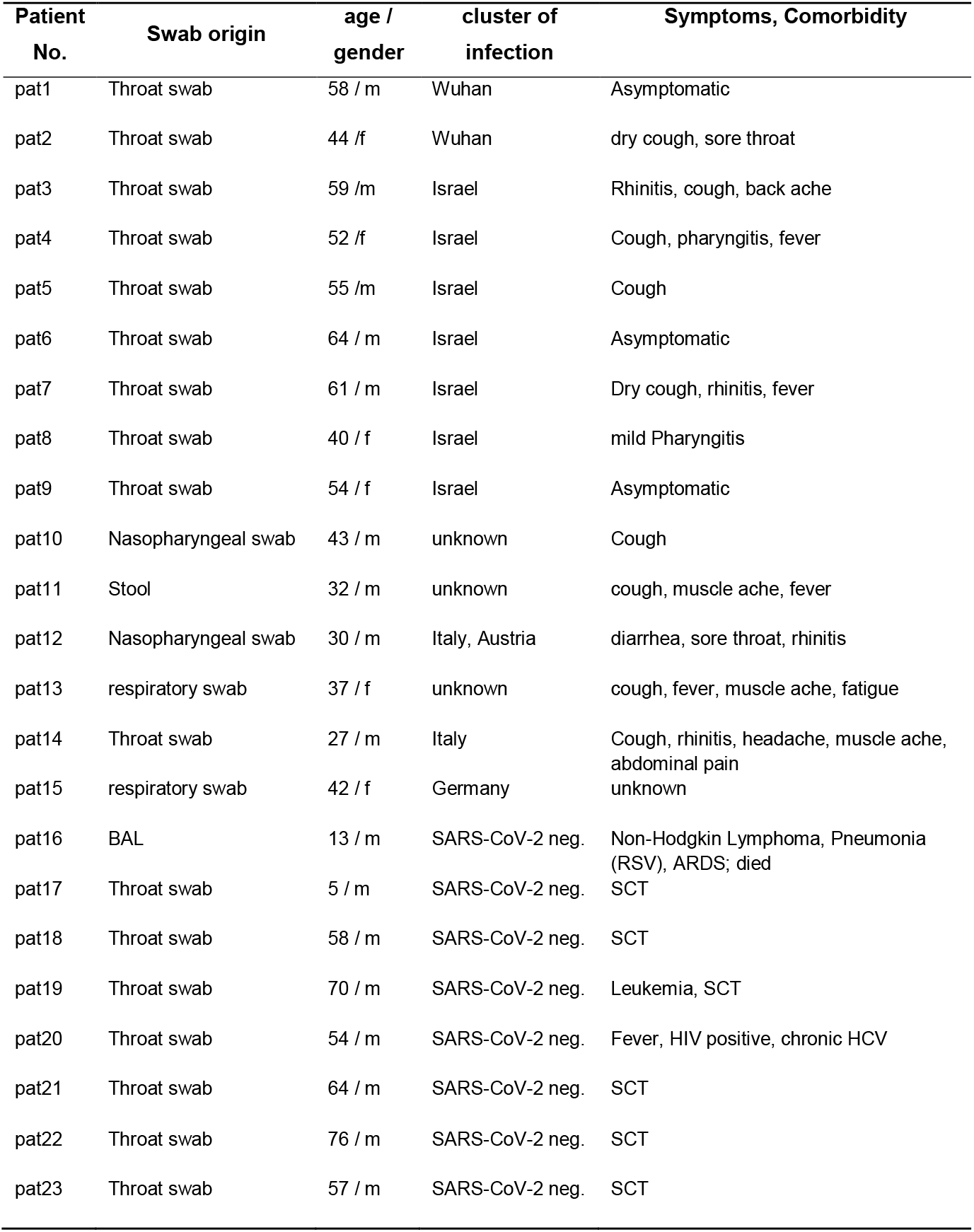
Compilation of patient samples used for validation of M-gene PCR in a clinical relevant setting.

